## Supplemental data for "Cohesin-independent STAG proteins interact with RNA and localise to R-loops to promote complex loading"

**Supplementary Figure Legends to accompany Porter, Li et al.**

**Figure S1. SA interacts with CTCF in the absence of cohesin.**

a) Immunoblot analysis of RAD21 levels in whole cell lysates from in RAD21$^{mAC}$ cells with and without CMV-OsTIR1 integration in the genome. Cells were untreated (-), treated with ethanol (EtOH) or with Auxin (IAA) for 4hrs. The IAA affect was also assessed with antibody to the AID tag. Tubulin and H3 serve as loading controls.

b) Representative confocal images of SA2 and CTCF IF in RAD21$^{mAC}$ cells treated with ethanol (EtOH) or Auxin (IAA) for 4hrs. Nuclei were counterstained with DAPI.

c) Chromatin coIP of SA1, SA2 and CTCF, together with Mock IgG controls in HeLa cells treated with siRNA to SMC3 for 48hrs.

d) GFP-TRAP in RAD21$^{mAC}$ cells immunoblotted for RAD21, SA1 and SA2. TRAP was performed with two concentrations of beads. Shown also is the flow-through material (post-TRAP) with % of residual proteins as determined by ImageJ densitometry.

e) Dual-color STORM images of SA2 (green) and CTCF (magenta) in EtOH and IAA-treated RAD21$^{mAC}$ cells. Representative full nuclei and zoomed nuclear areas are shown. Line denotes 2 microns and 200nm for full nuclei and zoomed areas respectively.

f) Nearest Neighbor Distance (NND) distribution plot of the distance difference between the experimental and random simulated data for SA1 (left) or SA2 (right) at CTCF localizations in EtOH (black) and IAA (red)-treated cells.

g) NND distribution plot of the distance between CTCF and SA1 (top left) or SA2 (bottom left) clusters in EtOH and IAA-treated cells. Experimental data are shown as continuous lines, random simulated data are displayed as dotted lines. Shown also are the NND distribution plots of the distance difference between the experimental and random simulated data for CTCF at SA1 (top right) or CTCF at SA2 (bottom right) localizations in EtOH (black) and IAA (red)-treated cells.

h) Pairwise comparisons of global CTCF ChIP-seq data (from merged biological replicates) compared to RAD21, SA1 and SA2 ChIP-seq in EtOH and IAA-treated RAD21$^{mAC}$ cells.

i) ChromHMM analysis of our ChIP-seq data as well as publicly available ChIP data in HCT116 cells as shown (Methods). Marks enriched within a given state (left) and enriched genomic features (right) are shown. We note that state 6 includes enrichments for ChIP data from RAD21 and SMC3 controls as well as SA1, SA2, CTCF in both control and IAA. This state is also enriched for active marks such as Polr2a, H3K4me3 and H3K27ac.

**Figure S2. Characterization of SA1 protein-protein interaction network in RAD21-depleted cells.**

a) SlimSearch motifs for the shown 'CES' binding proteins. The CES-binding motif is capitalised and the F/YXF-motif within is highlighted in red.

b) Replicate chromatin coIP of SA1, SA2 and IgG with predicted CES-binding proteins in RAD21$^{mAC}$ cells treated with EtOH or IAA for 4hrs. Input represents 1.25% of the material used for immunoprecipitation.

Enriched biological processes for the c) SA1 interactome and d) SA1$^{\Delta Coh}$ interactome compared to the whole genome.

e) Full SA1$^{\Delta Coh}$ interaction network of protein–protein interactions identified in RAD21$^{mAC}$ cells using STRING. Colours and annotations as in Fig. 2c.

**Figure S3. SA proteins bind to RNA and localise to R-loops.**
CLIP for a) SA1 and b) SA2 treated with various controls as shown. Autoradiograms of crosslinked $^{32}$P-labelled RNA are shown at the top and the corresponding immunoblots, below.

c) CLIP for SA1, SA2 and IgG control in cells treated with siRNAs to scramble control (siscr), SA1 (siSA1) or SA2 (siSA2). $^{32}$P-labelled RNA and the corresponding immunoblots are shown as above. Input material (WCL) is included on the blots. *NB* the loss of the RNA signal upon siRNA KD.

d) Immunoblot analysis of RAD21, SA1 and SA2 levels in whole cell lysates (WCL) from RAD21$^{mAC}$ cells treated with EtOH or IAA. Actin serves as a loading control.

e) CLIP for SA1, SA2 and IgG control in EtOH (-) or IAA-treated (+) samples from d). $^{32}$P-labelled RNA and the corresponding immunoblots are shown as above. *NB* SA1 and SA2 samples were loaded to show proportional amounts of the proteins in the IAA condition.

f) Representative confocal images of S9.6 and AQR (leftmost panel), SA1 (middle panel) and SA2 (rightmost panel) IF in RAD21$^{mAC}$ cells treated with scramble control siRNA (si Con) or siRNA to the protein of interest. Nuclear outlines (white) are derived from DAPI counterstain. Mean fluorescence Intensity (MFI) shown below images. Data are from three biological replicates with >50 cells counted/condition. Quantifications and statistical analysis were done as previously stated.

g) Imaris quantification of (left) mClover or (right) S9.6 signal by IF in RAD21$^{mAC}$ cells treated with EtOH or IAA for 4hr. Data are from three biological replicates with >50 cells counted/condition). *NB* We detected a change in S9.6 signal upon IAA treatment by IF, however this did not reach statistical significance. Quantifications and statistical analysis were done as indicated previously.

h) Replicate chromatin coIP of S9.6 and IgG in RAD21$^{mAC}$ cells treated with RNase H enzyme (RNH) and immunoblotted with antibodies representing known R-loop proteins and both SA1 and SA2. Input represents 1.25% of the material used for immunoprecipitation. Bottom, S9.6 dot blot of lysates used in coIP.

i) Representative S9.6 dot blot of two dilutions of chromatin samples used in the S9.6 coIPs and treated with enzymes as shown. A positive control for the digestion of RNA:DNA hybrids was included with a high concentration and temperature global RNase A digestion sample.

j) Summary plots showing mean ChIP-seq read density across the regions from Fig 3h including the SA1 ChIP-seq data.

k) ChromHMM analysis as in Fig S1i and now including our DRIP-seq data from HCT116 cells. The 'R-loop_SA overlap' and 'R-loop_SA adjacent' sites are marked in red. Marks enriched within a given state (left) and enriched genomic features (right) are shown. We note that 'R-loop_SA overlap' sites enrich genes and active marks such as Polr2a, H3K4me3 and H3K27ac in states 2/3/12 while the 'R-loop_SA adjacent' sites cluster separately with repressive chromatin marks such as Lamin B1 LADs in states 7/8.

**Figure S4. SA proteins contribute to cohesin loading**
a) mClover MFI shown as in Fig 4b and including a IAA-washoff sample treated with auxinole (an Auxin inhibitor) which leads to increased recovery of mClover signal in cells.

b) Schematic of experimental set-up. RAD21$^{mAC}$ cells expressing mClover (green cells in dishes) were treated with scramble siRNAs or siRNA to NIPBL. Prior to collection, cells were cultured in EtOH or IAA for 4hrs to degrade RAD21 (0 h timepoints). The EtOH or IAA treatment was washed-off and the cells were left to recover for 4hrs (4h timepoints). Chromatin fractions were prepared from all samples and used in immunoblot analysis.

c) Full immunoblot analysis of the data shown in main Fig. 4d. Chromatin-bound RAD21, NIPBL and MAU2 levels in RAD21$^{mAC}$ cells treated according to the schematic shown in Fig. S4b as well as a longer timepoint of 're-loading' (22hr, which we found to be too stressful to cells). H3 was used as a loading control.

d) Quantification of the RAD21 fold change relative to siCon samples at the 0h timepoint in siCon, siNIPBL, siSA and siNIPBL+siSA from immunoblot analysis of chromatin-bound RAD21 in main Fig. 4f. Asterisks indicate a statistically significant difference as assessed using 2-tailed T-test. P-values as before. Data is from 5 biological replicates.

e) Immunoblot analysis of chromatin-bound RAD21, SA1, SA2, and H3 levels in RAD21mAC cells treated according to the schematic shown in Fig. S4b and including samples treated with siRNA to SA1, SA2 and SA1 and SA2 together. H3 was used as a loading control.

**Figure S5. A basic exon in SA2 influences RBP stability.**

a) Percent Spliced In (PSI) calculations for SA1 exon 31 and SA2 exon 32 based on VAST-Tools analysis of RNA-seq in three separate datasets and the average (AVG) reported in the manuscript. Reads crossing exon junctions from each sample are also shown.

b) PSI distributions for SA1 exon 31 and SA2 exon 32 based on VAST-Tools analysis of RNA-seq from mouse ESCs and neural progenitors (NPC).

c) Peptide profile plots from YFP-SA2 overexpression CLIP-MS experiment from Fig 5e. Shown are the Log2 peptide intensities for each protein in each biological replicate IP from GFP-TRAP alone (mock IP), YFP-SA2-e32 and YFP-SA2-FL. Core cohesin components, YFP and STAG2 are robustly IP'd in all samples.

d) Enriched biological processes for the YFP-SA2 interactome compared to published SA2 mass spec study in Kim et al. NB. RNA-binding proteins are enriched in both.

**Supplementary Figure 1.**

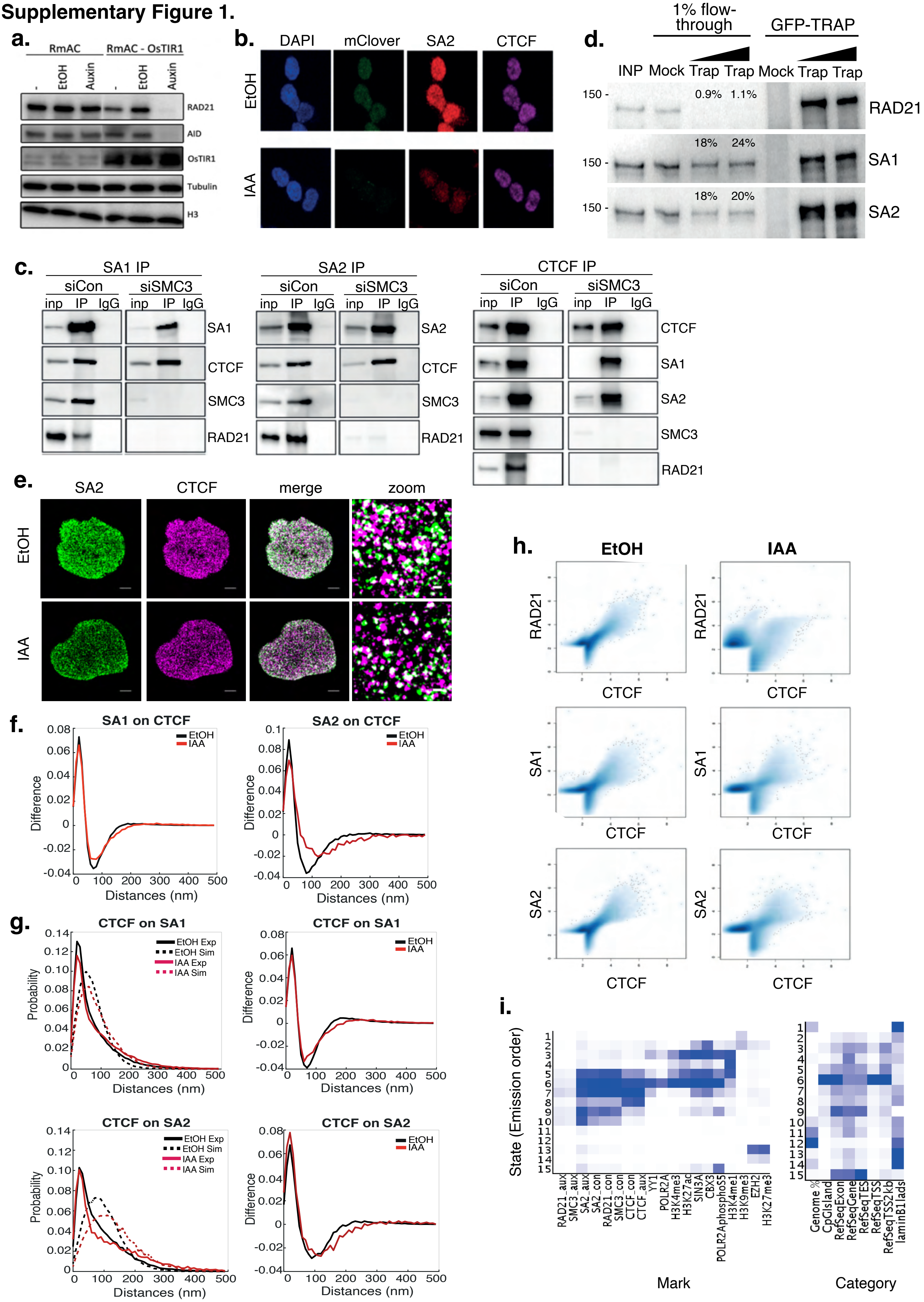

#### Supplementary Figure 2.

**a.**

| Name | Protein Accession | Protein Name | Motif |
|---|---|---|---|
| CTCF | P49711 | Transcriptional repressor CTCF | dvdvsVYDFEEEqqegl |
| MCM3 | P25205 | DNA replication licensing factor MCM3 | gdsydPYDFSDTEeempq |
| HNRNPUL2 | Q1KMD3 | Heterogeneous nuclear ribonucleoprotein U-like protein 2 | ehgraYYEFREEayhsr |
| CHD6 | Q8TD26 | Chromodomain-helicase-DNA-binding protein 6 | qkhrrPYEFEVErdaka |
| ESYT2 | A0FGR8 | Extended synaptotagmin-2 | sqrrsAFGFDDgnfpg |

**b.**

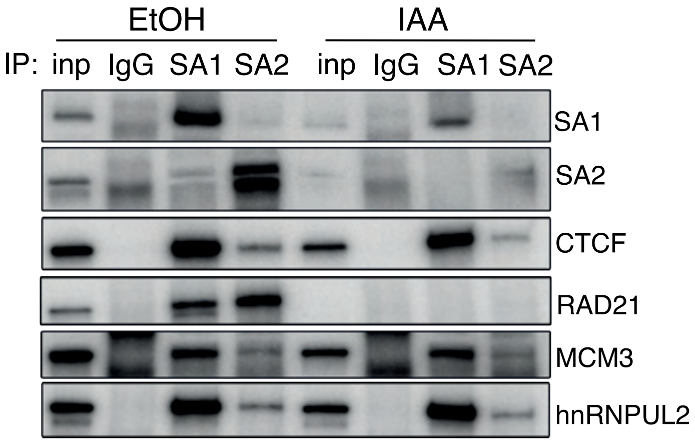


**c.**

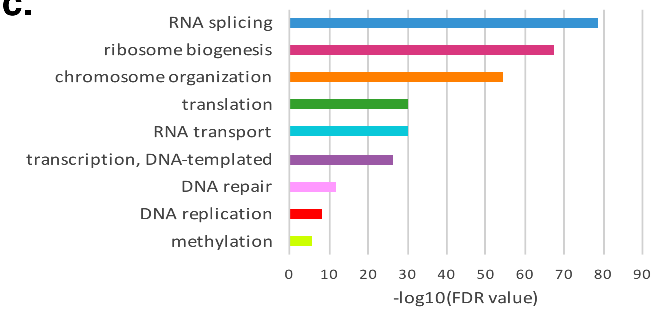


**d.**

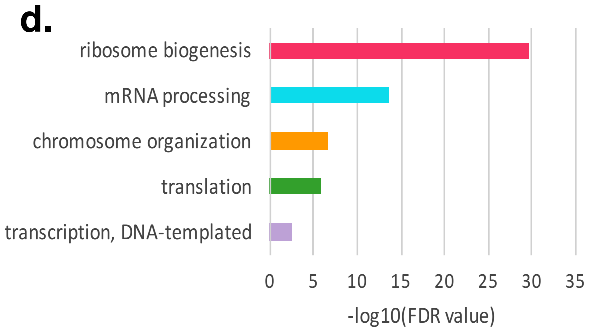


**e.**

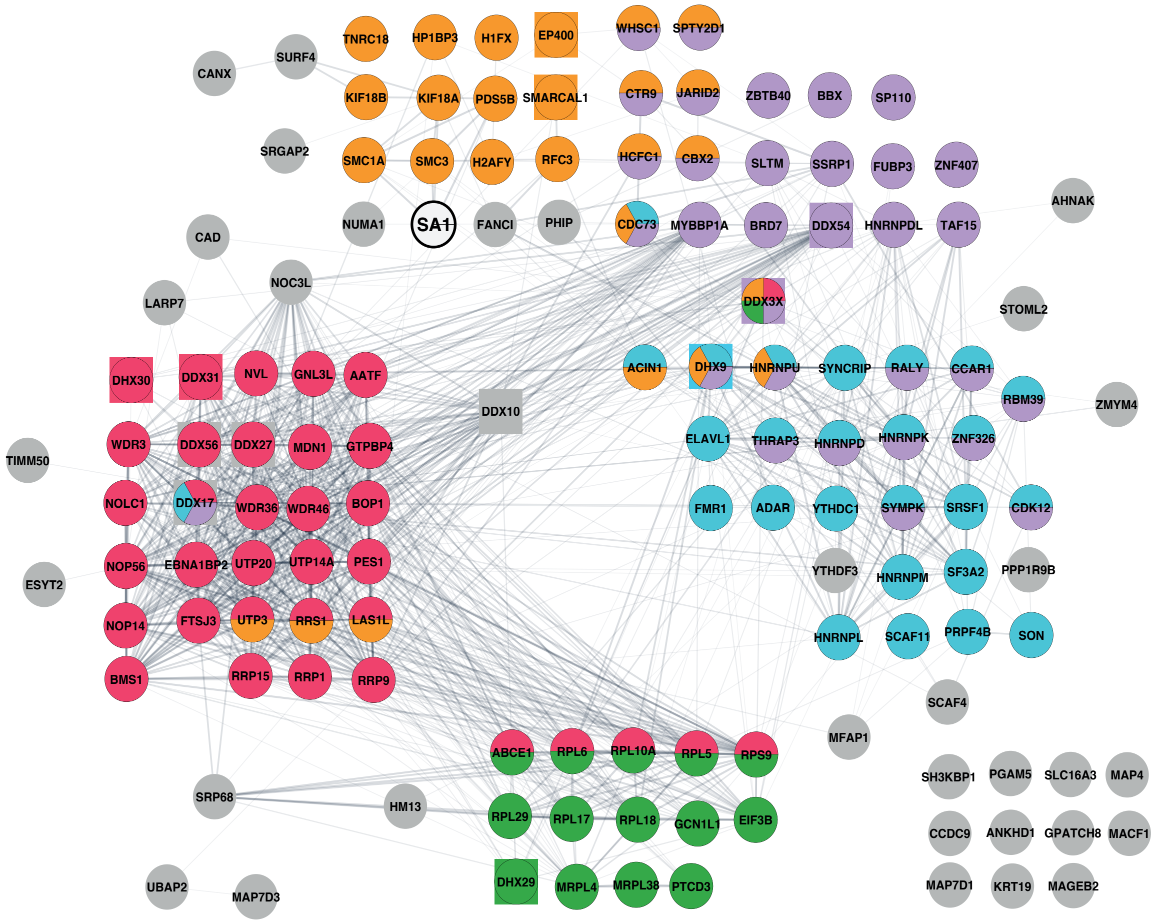

#### Supplementary Figure 3.

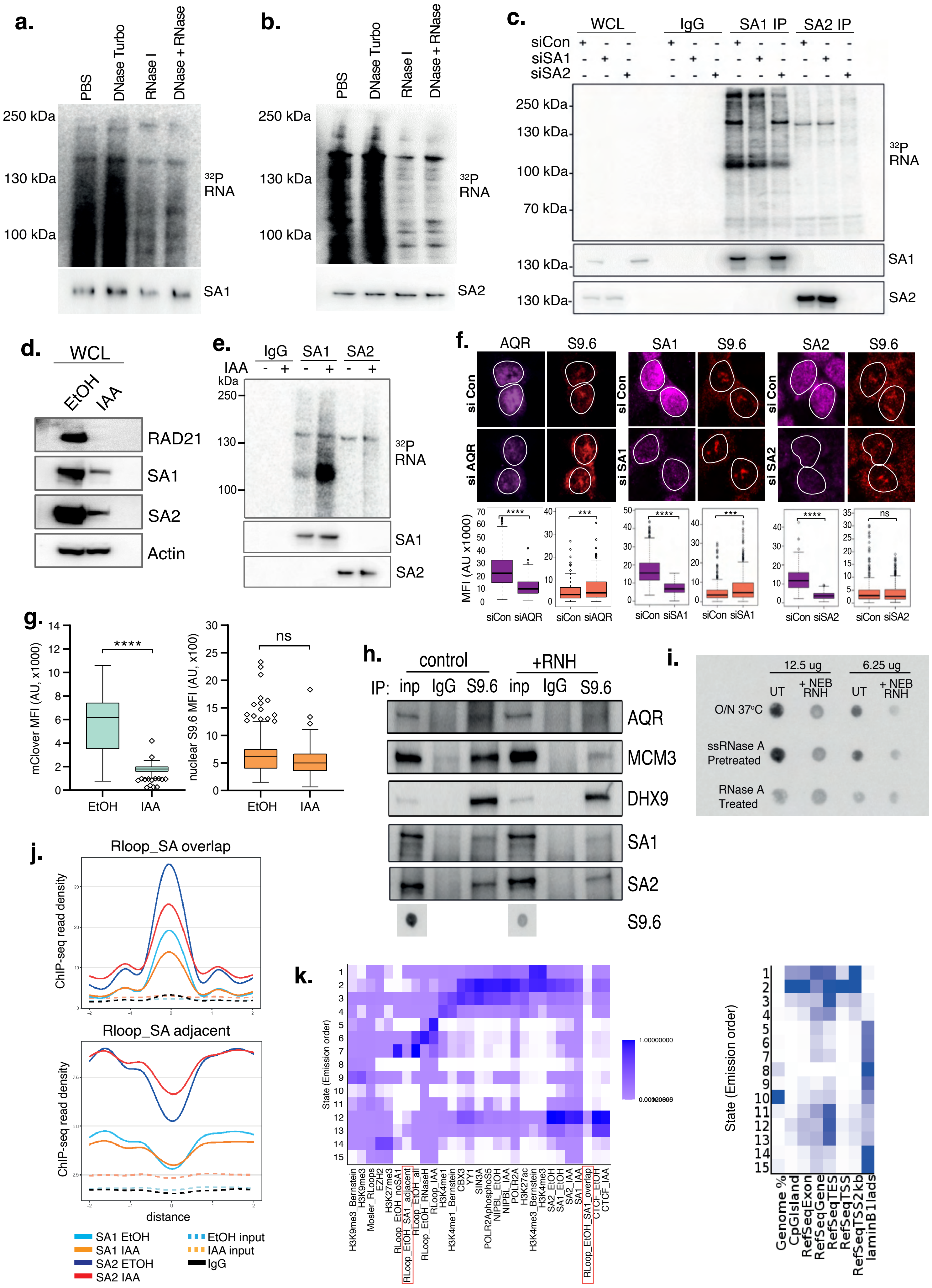

### Supplementary Figure 4.

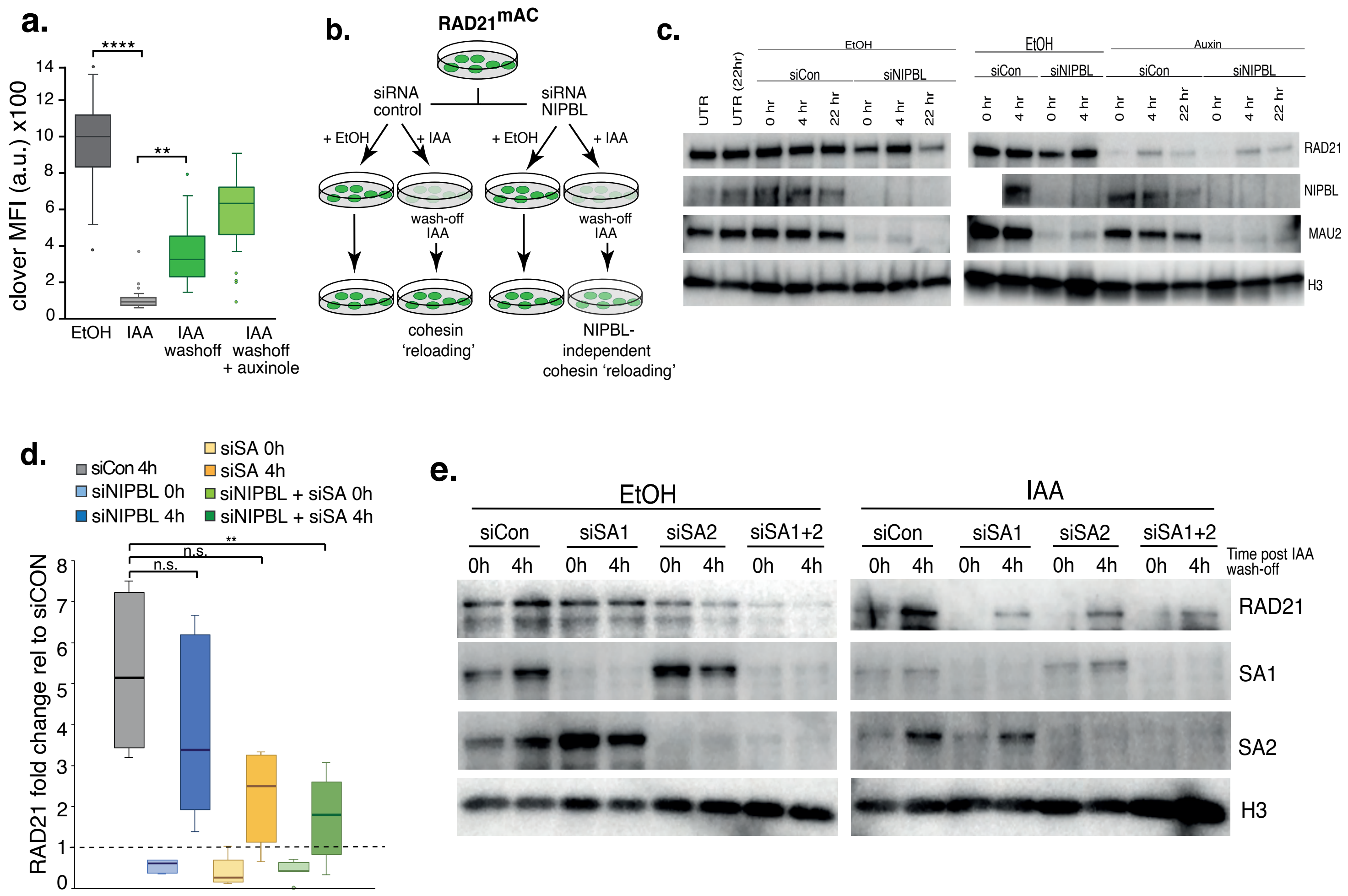

### Supplementary Figure 5.

**a.**

| Dataset | SA2 e32 PSI | Reads across junction | SA1 e31 PSI | Reads across junction |
|---|---|---|---|---|
| ENCODE HCT | 21.92 | 202.98 | 98.99 | 588 |
| Zuo HCT | 19.18 | 94.75 | 97.91 | 278.06 |
| ENCODE HeLa | 19.95 | 90.57 | 96.36 | 156.1 |
| **AVG** | **20.35** | | **97.75** | |

**b.**

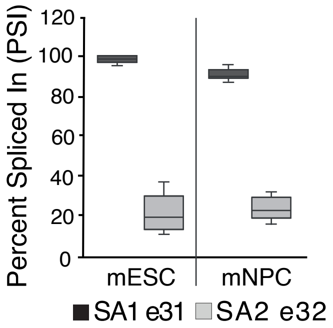


**c.**

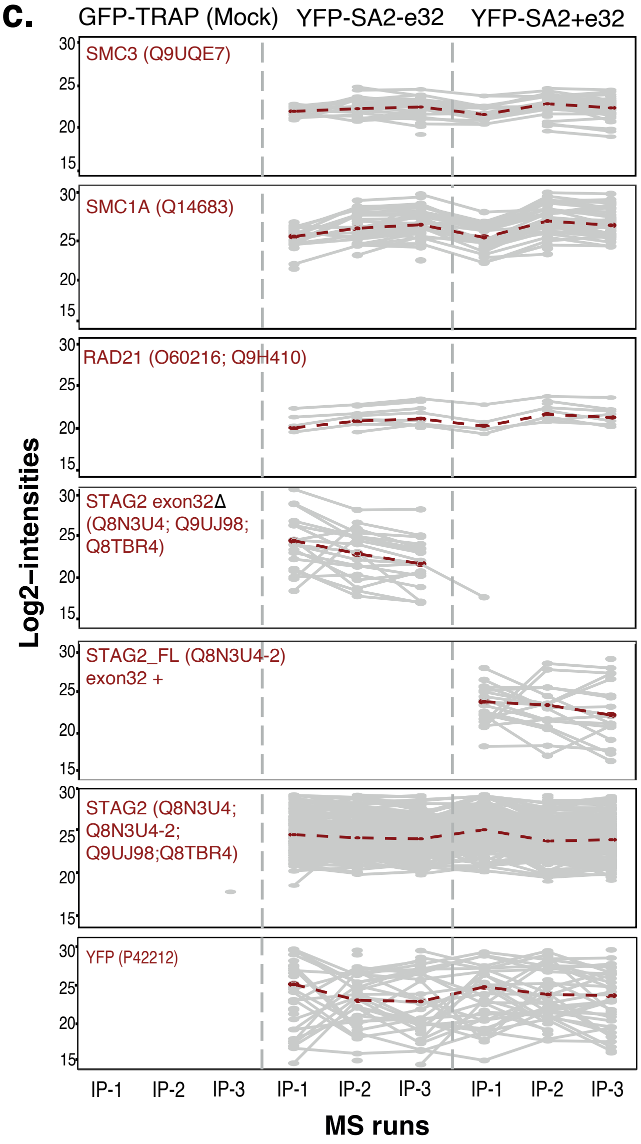


**d.**

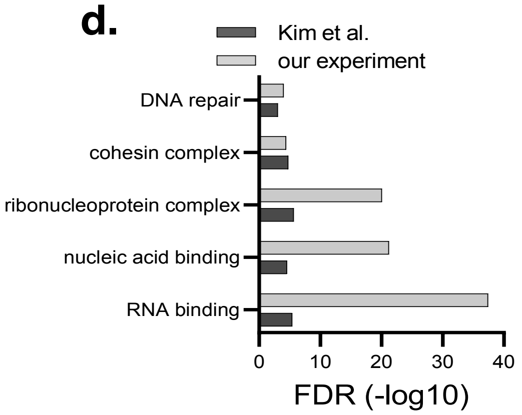
